# TSC2 regulates tumor susceptibility to TRAIL-mediated T cell killing by orchestrating mTOR signaling

**DOI:** 10.1101/2022.05.09.491124

**Authors:** Chun-Pu Lin, Joleen J.H. Traets, David W. Vredevoogd, Nils L. Visser, Daniel S. Peeper

**Affiliations:** Division of Molecular Oncology and Immunology, Oncode Institute, The Netherlands Cancer Institute, Plesmanlaan 121, 1066 CX, Amsterdam, the Netherlands; Division of Tumor Biology and Immunology, The Netherlands Cancer Institute, Plesmanlaan 121, 1066 CX, Amsterdam, the Netherlands

**Keywords:** mTOR, T cell sensitivity, TRAIL, TSC2, tumor cells

## Abstract

Resistance to cancer immunotherapy continues to impair common clinical benefit. Here, we uncover an important role for Tuberous Sclerosis Complex 2 (TSC2) in determining tumor susceptibility to cytotoxic T lymphocytes (CTL) killing, both *in vitro* and *in vivo*. *TSC2*-depleted tumor cells showed disrupted mTOR regulation upon CTL attack, which was associated with enhanced cell death. Tumor cells adapted to CTL attack by shifting their mTOR signaling balance toward increased mTORC2 activity to circumvent apoptosis and necroptosis. TSC2 critically protected tumor cells as its ablation strongly augmented tumor cell sensitivity to CTL attack. Mechanistically, TSC2 inactivation caused elevation of TRAIL receptors expression, cooperating with mTORC1-S6 signaling to induce tumor cell death. Clinically, we found a negative correlation between *TSC2* expression and TRAIL signaling in TCGA patient cohorts. Moreover, a lower TSC2 immune response signature was observed in melanomas from patients responding to immune checkpoint blockades. Our study uncovers a pivotal role for TSC2 in the cancer immune response by governing crosstalk between TSC2-mTOR and TRAIL signaling, aiding future therapeutic exploration of this pathway in immuno-oncology.

## Introduction

Immunotherapy, particularly immune checkpoint blockade (ICB), has transformed cancer patient care in recent years. The blockade of inhibitory immune checkpoints, such as programmed cell death 1 (PD-1) and cytotoxic T lymphocyte-associated protein 4 (CTLA-4), unleashes a potent anti-tumor response with cytotoxic T cells (CTL) for a growing number of patients (Hodi *et al*, 2010; Larkin *et al*, 2015; Wolchok *et al*, 2017; Schadendorf *et al*, 2017; Larkin *et al*, 2019).

CTL are activated when their T cell receptors (TCRs) encounter matching antigens presented by antigen-presenting cells (APC) or tumor cells. This can result in the elimination of target cells by the secretion of cytotoxic molecules, death ligands and cytokines triggering cell death signaling (Martínez-Lostao *et al*, 2015; Russell & Ley, 2002; Farhood *et al*, 2019). Owing to the critical role of CTL in cancer immunotherapy, a rapidly increasing number of immunotherapeutic studies have been launched focusing on reversing T cell dysfunction to improve treatment outcome (Zarour, 2016; Wherry & Kurachi, 2015; Jiang *et al*, 2018; Thommen & Schumacher, 2018). However, tumor-intrinsic mechanisms, too, often contribute to the escape of immune surveillance, commonly impairing durable responses to ICB (Zaretsky *et al*, 2016; Spranger *et al*, 2015; Gao *et al*, 2016b; Litchfield *et al*, 2021; Sharma *et al*, 2017).

Among various resistance mechanisms, IFNγ signaling plays a crucial role in determining tumor sensitivity to T cells. For example, tumors with specific deficiencies in IFNγ signaling can be more resistant to immune checkpoint therapy (Zaretsky *et al*, 2016; Shin *et al*, 2017; Gao *et al*, 2016a). Therefore, we previously set out to identify IFNγ signaling-independent tumor determinants of T cell sensitivity. Specifically, we performed an unbiased genome-wide CRISPR-Cas9 knockout screen in IFNγ receptor-deficient (*IFNGR1*-KO) melanoma cells under cytotoxic T cell attack, uncovering an important role of TRAF2 in determining tumor sensitivity to T cell elimination in an IFNγ-independent tumor landscape (Vredevoogd *et al*, 2019). In the current study, we re-analyzed the results of this screen and describe a novel regulator of tumor cell sensitivity to T cell killing, both *in vitro* and *in vivo*. We also investigated its mechanism of action, and clinically corroborated the results.

## Results

### Whole-genome CRISPR-Cas9 knockout screen identifies TSC2 as a negative regulator of tumor sensitivity to CTL killing

To identify critical factors protecting tumor cells against CTLs in addition to TRAF2 (Vredevoogd *et al*, 2019), we re-analyzed the data from our previous genome-wide CRISPR-Cas9 knockout screen (Fig. 1A). The screen was performed in a human HLA-A*02:01^+^/MART1^+^ Interferon Gamma Receptor 1-knockout (*IFNGR1*-KO) D10 melanoma cell line that was challenged with healthy donor CD8 T cells, which were retrovirally transduced with a MART-1-reactive T cell receptor (Gomez-Eerland *et al*, 2014). Tumor cells expressing sgRNAs targeting Tuberous Sclerosis Complex Subunit 1 or 2 (*TSC1* and *TSC2*) were significantly depleted from tumor cells under attack by MART-1-specific T cells, indicating that these genes contribute to tumor cell-intrinsic susceptibility to T cells (Fig. 1B). TSC1 and TSC2 are known to be crucial regulators of many biological processes by forming a complex that negatively regulates mTORC1 via the GTPase activation property of TSC2 towards RheB (Inoki *et al*, 2003a; Zhang *et al*, 2003; Tee *et al*, 2003; Garami *et al*, 2003). Their dysregulation contributes to tumor development (Menon & Manning, 2009; Jiang *et al*, 2005; Adachi *et al*, 2003; Xu *et al*, 2009). This correlates with elevated mTORC1 signaling (Inoki *et al*, 2002; Potter *et al*, 2002), which in turn leads to increased cell metabolism and biosynthesis while inhibiting autophagy, ultimately resulting in enhanced cell growth (Düvel *et al*, 2010; He *et al*, 2018; Valvezan *et al*, 2017; Hosokawa *et al*, 2009; Kim *et al*, 2011, 2002; Inoki *et al*, 2003b, 2006; Liu & Sabatini, 2020). However, how TSC1 and TSC2 regulate tumor vulnerability to T cell toxicity has not yet been addressed to our knowledge.

**Figure 1:**
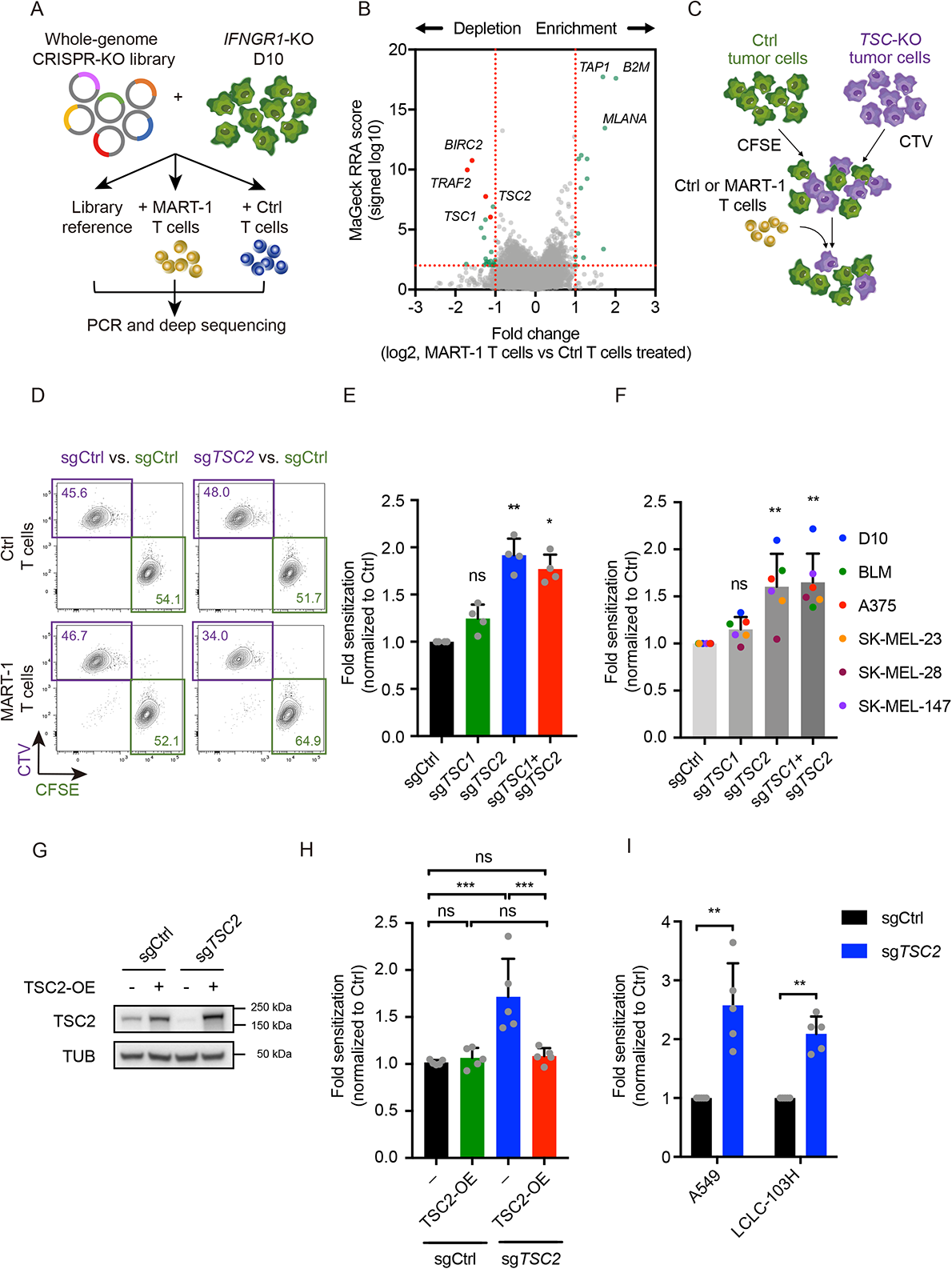
Whole-genome CRISPR-Cas9 knockout screen identifies TSC2 as a negative regulator of tumor sensitivity to CTL killing. A) Schematic outline of *in vitro* genome-wide CRISPR-Cas9 knockout screen in IFNγ receptor-deficient (*IFNGR1*-KO) D10 melanoma cells under CD8 T cell challenge (Vredevoogd *et al*, 2019). B) Volcano plot of depleted and enriched sgRNAs in tumor cells treated with MART-1 T cells versus Ctrl T cells. C) Schematic illustration of *in vitro* competition assay. D) Representative flow cytometry plot of *in vitro* competition assay. E) *In vitro* tumor T cell co-culture competition assay in *IFNGR1*-KO D10 cells containing sgRNAs targeting *TSC1, TSC2* or both. Fold sensitization is calculated by the ratio change of remaining cells expressing sgCtrl versus knockout cells, relative to Ctrl T cell groups. Statistics was done by Kruskal-Wallis test with Dunn’s post hoc test. Error bars indicate SD of four independent experiments with different T cell donors (n = 4). * P < 0.05; ** P < 0.01; *** P < 0.001; **** P < 0.0001. F) *In vitro* tumor T cell co-culture competition assay in multiple melanoma cell lines containing indicated sgRNAs. Statistics was done by Kruskal-Wallis test with Dunn’s post hoc test. Error bars indicate SD of six independent experiments with different cell lines (n = 6). * P < 0.05; ** P < 0.01; *** P < 0.001; **** P < 0.0001. G) Western blot analysis of *TSC2* expression in cells used in (H). H) Competition assay performed in sgCtrl- and sg*TSC2*-expressing D10 cells, and cells reconstituted with *TSC2* expression. Statistical analysis was performed with a one-way ANOVA, followed by a Tukey post-hoc test. Error bars indicate SD of five independent experiments with different T cell donors (n = 5). * P < 0.05; ** P < 0.01; *** P < 0.001; **** P < 0.0001. I) *In vitro* competition assay of sgCtrl and sg*Tsc2*-expressing lung cancer cell lines co-cultured with MART-1 T cells normalized to the Ctrl T cell-treated group. Statistics was done by Mann–Whitney test. Error bars indicate SD of five independent experiments with different T cell donors (n = 5). * P < 0.05; ** P < 0.01; *** P < 0.001; **** P < 0.0001.

TSC1 and TSC2 act in a complex to inhibit mTORC1 activity. To validate these screen hits and assess their contributions to the sensitization to cytotoxic T cells, we first generated either single *TSC1*, *TSC2* or *TSC1/TSC2* double knockout (KO) tumor cell lines. Because the screen was done in a *IFNGR1*-KO background, this was also the first setting we tested here. After differentially labelling non-targeting sgRNA expressing cells (Ctrl) and TSC-KO cells with CFSE and CTV, respectively, the control and TSC-KO cells were mixed at a 1:1 ratio. This tumor cell mix was subsequently co-cultured with either non-tumor-reactive (Ctrl) or MART-1-reactive CD8^+^ T cells for 3 days. T cell sensitivity was assessed by the ratio of Ctrl to TSC-KO tumor cells that survived at the end of co-culture (Fig. 1C). Flow cytometry analysis showed that loss of *TSC2* alone or *TSC1/TSC2* double-KO significantly sensitized *IFNGR1*-KO melanoma cells to T cell killing to a similar extent, whereas TSC1-KO alone showed a similar trend but was statistically insignificant (Fig. 1D, E). These results indicate that TSC2 is the essential component of the TSC1-TSC2 complex that limits sensitivity to T cell attack in *IFNGR1*-deficient tumor cells. This aligns with the notion that TSC1 acts as a stabilizer of TSC2 instead of having a direct GTPase activating function (Chong-Kopera *et al*, 2006; Benvenuto *et al*, 2000), limiting its dominancy in the regulation.

To investigate whether these observations are dependent on the absence IFNγ signaling, we used IFNγ signaling-proficient cells. To avoid cell line bias, we used a cell line panel and performed the same competition assay, producing similar results (Fig. 1F). Together, these results suggest a general and rate-limiting role of TSC2 in regulating tumor cells sensitivity to cytotoxic T cells.

TSC2 accounts for the main functional activity in the TSC complex by acting as a GTPase-activating protein (GAP) for the small GTPase Rheb, a direct mTORC1 activator (Castro *et al*, 2003). Since the effect of TSC2 inactivation was comparable to the combined knockout of both *TSC1* and *TSC2*, we decided to focus on the influence of *TSC2* depletion in the regulation of tumor cells to T cell attack. The *TSC2*-depletion-induced T cell sensitization was validated with multiple *TSC2*-targeting sgRNAs **(Fig. EV1A and EV1B).** Further excluding sgRNA off-target effects, we reintroduced *TSC2* cDNA into *TSC2*-knockout melanoma cells (Fig. 1G). This led to a rescue of the enhanced tumor sensitivity to T cell killing, demonstrating the specificity of this phenotype to *TSC2* inactivation (Fig. 1H). We next validated the T cell-sensitizing effect caused by *TSC2*-depletion in multiple human lung cancer cell lines (Fig. 1I), as well as in B16 murine melanoma cells **(Fig. EV1C and EV1D).** Thus, TSC2 ablation causes a general and robust sensitivity to T cell cytotoxicity across different cell types of mouse and human origin.

### *TSC2* ablation sensitizes melanoma cells to CTL killing in an *in vivo* tumor ACT model

To assess the translational value of our *in vitro* findings, we next tested whether TSC2 deficiency can sensitize tumors to T cell-mediated tumor elimination *in vivo* as well. We performed an *in vivo* competition assay with a similar experimental setup to our *in vitro* competition assay, where *TSC2*-KO and Ctrl melanoma cells expressing either mCherry or eGFP respectively, were mixed at a 1:1 ratio. Next, they were subcutaneously transplanted into NOD severe combined immunodeficiency (SCID) gamma/B2m-deficient (NSG) mice (Fig. 2A). Adoptive cell transfer (ACT) was performed with either Ctrl T cells or MART-1 T cells, after tumors had established. Mice treated with Ctrl T cells showed steady tumor outgrowth. In contrast, significant tumor control was observed in mice receiving 1D3 T cell transfer, indicating the presence of an antigen-specific antitumor effect of ACT (Fig. 2B). Tumor subpopulations that had survived the ACT were harvested and analyzed by flow cytometry. Compared to the control T cell-treated tumors, tumors treated with MART-1 T cells showed a depletion of *TSC2*-KO cells (Fig. 2C). In line with our *in vitro* results, the quantification of *in vivo* competition assay showed that tumor cells with *TSC2*-KO were considerably more vulnerable to CD8^+^ T cells also *in vivo* (Fig. 2D). The same effect was observed in an *in vivo* competition assay with reversed color labeling, excluding a label-specific effect **(Fig. EV2A-EV2C)**. These results strengthen the notion that TSC2 serves a critical determinant of tumor sensitivity to cytotoxic T cell pressure.

**Figure 2:**
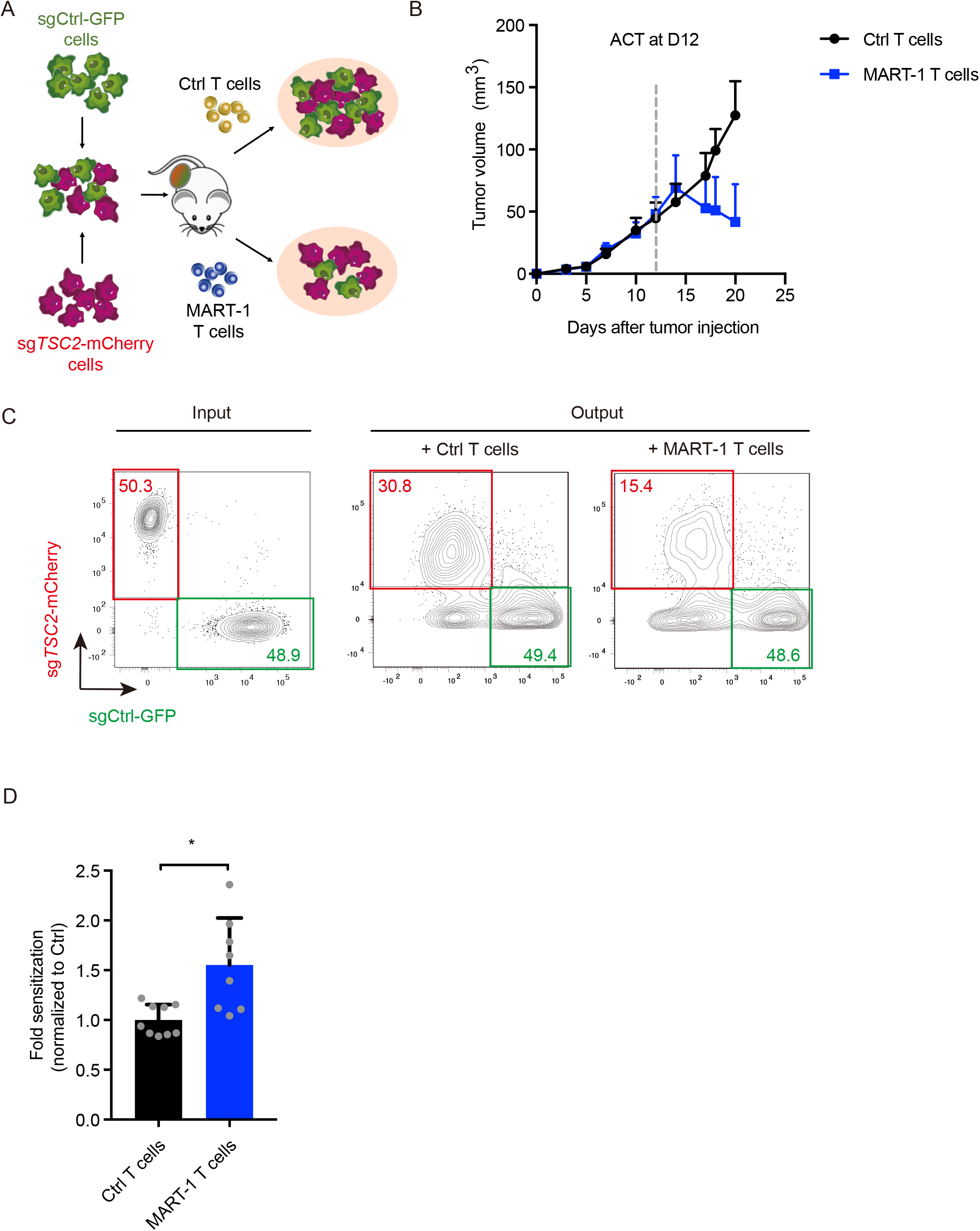
TSC2 ablation sensitizes melanoma cells to CTL killing in an *in vivo* tumor ACT model. A) Schematic outline of *in vivo* competition assay. B) Tumor growth upon adoptive cell transfer with Ctrl (n = 9) or MART-1 (n = 8) T cells. Data points represent average tumor volume, and error bars represent SEM. C) Flow cytometry plot of both tumor mix input and representative output of the *in vivo* competition assay. D) Quantification of the *in vivo* competition assay (output) at end point by flow cytometry analysis. Error bars indicate SD. Statistical analysis was performed by Mann–Whitney test. * P < 0.05; ** P < 0.01; *** P < 0.001; **** P < 0.0001.

### *TSC2* depletion-induced deregulation of mTOR signaling increases tumor cell death by CTL attack

While TSC1-TSC2 complex is known as a pivotal inhibitor of mTORC1 activity, it can also physically interact with mTORC2 (Huang *et al*, 2008; Frias *et al*, 2006), which in turn contributes to the phosphorylation of Akt, promoting cell proliferation and survival. Moreover, chronically activated mTORC1 signaling can attenuate PI3K/Akt downstream signaling, including cell survival and proliferation, serving as a negative feedback mechanism (Harrington *et al*, 2004; Shah *et al*, 2004). In agreement with the reported phenotype in melanocytes (Cao *et al*, 2017), we noted that TSC2 ablation triggered an induction of mTORC1 signaling in multiple melanoma and lung cancer cell lines, indicated by the enhanced phosphorylation of ribosomal protein S6 at phospho-sites Ser235/236 and Ser240/244 (Fig. 3A). This was accompanied by a more heterogenous downregulation of mTORC2 signaling, as indicated by the suppression of phosphorylated Akt levels in most *TSC2*-KO cell lines. However, we noted a consistently higher mTORC1:mTORC2 signaling ratio as judged by the relative phosphorylation level of ribosomal protein S6 and Akt in *TSC2*-KO cells, indicating an mTORC1-skewing effect by *TSC2*-depletion (Fig. 3B). A similar effect was observed in TSC1 or *TSC1*/*TSC2* double knockout cells. Again supporting the dominant role of TSC2 in regulating mTOR signaling, the effect was only minor in TSC1-KO alone **(Fig. EV3A)**.

**Figure 3:**
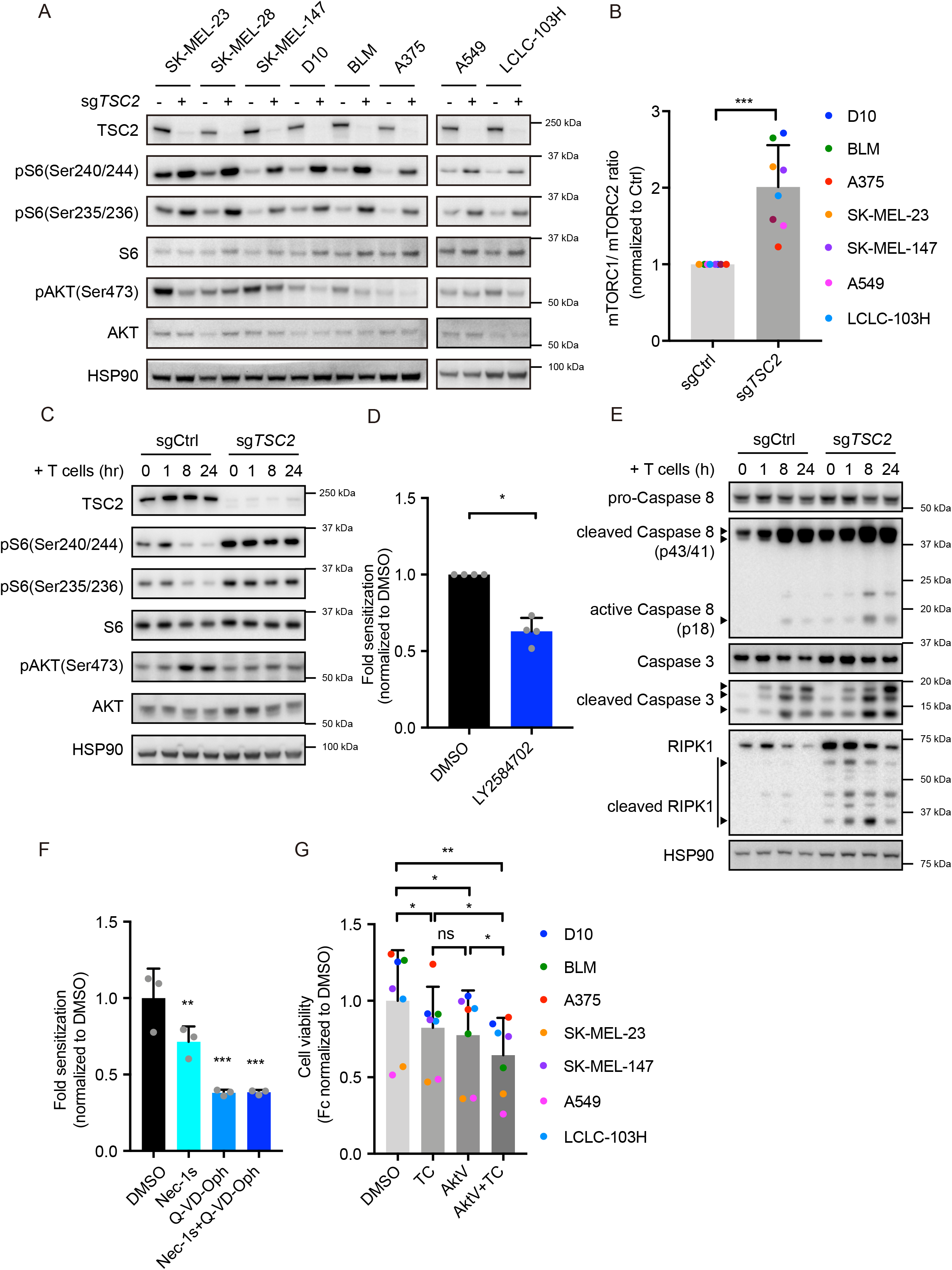
*TSC2* depletion-induced deregulation of mTOR signaling increases tumor cell death by CTL attack. A) Western blot analysis of baseline mTOR signaling in sgCtrl- or sg*Tsc2*-expressing melanoma and lung cancer cell lines. Representative plot of 2-6 independent experiments. All blots shown in this panel were run in parallel. B) Western blot quantification for mTORC1/mTORC2 signaling ratio from (A). mTORC1 signaling level was calculated by normalizing phosphorylated ribosomal protein S6 (Ser240/244) to total S6 expression; and mTORC2 signaling level was calculated by normalizing phosphorylated Akt (Ser473) to total Akt expression. Statistical analysis was performed by Mann–Whitney test. Error bars indicate SD of seven independent experiments with different cell lines (n = 7). * P < 0.05; ** P < 0.01; *** P < 0.001; **** P < 0.0001. C) Western blot analysis of mTOR signaling in sgCtrl- or sg*TSC2*-expressing D10 melanoma cells upon MART-1 T cell challenge for indicated time. Representative of 3 experiments with different T cell donors (n = 3). D) *In vitro* competition assay of sgCtrl- and sg*TSC2*-expressing D10 cells co-cultured with MART-1 T cells in the presence of LY2584702 (5uM), normalized to DMSO-treated groups. Statistical analysis was performed by Mann–Whitney test. Error bars indicate SD of four independent experiments with different T cell donors (n = 4). * P < 0.05; ** P < 0.01; *** P < 0.001; **** P < 0.0001. E) Western blot analysis of sgCtrl- or sg*TSC2*-expressing D10 melanoma cells upon MART-1 T cell challenge for indicated time. Representative of 3 experiments with different T cell donors (n = 3). F) *In vitro* competition assay of sgCtrl- and sg*Tsc2*-expressing D10 cells co-cultured with MART-1 T cells in the presence of necroptosis (Nec1-s, 20uM) or apoptosis inhibitors (Q-VD-Oph, 50uM), normalized to DMSO-treated groups. Statistical analysis was performed by one-way ANOVA with Holm-Sidak’s multiple comparisons test. Error bars indicate SD of three independent experiments with different T cell donors (n = 3). * P < 0.05; ** P < 0.01; *** P < 0.001; **** P < 0.0001. G) Cell viability analysis by CellTiter-Blue assay of T cell sensitivity in multiple melanoma and lung cancer cell lines in the presence of Tautomycin (TC, 150nM), Akt Inhibitor V (AktV, 1.5uM), or the combination (TC, 150nM^+^ AktV, 1.5uM) compared to DMSO-treated group. Statistical analysis was performed with a one-way ANOVA, followed by a Tukey post-hoc test. Error bars indicate SD of independent experiments with seven different cell lines. Each data point represent the average mean from three independent experiments with different T cell donors (n = 3). * P < 0.05; ** P < 0.01; *** P < 0.001; **** P < 0.0001.

To better understand the dynamics mTORC1 and mTORC2 signaling in response to cytotoxic T cells, particularly how they are affected by the absence of TSC2, we challenged either Ctrl or *TSC2*-KO tumor cells with MART-1 T cells and assessed the effects on mTOR signaling. In parental melanoma tumor cells challenged with MART-1 T cells, we observed a reduction in phosphorylated ribosomal protein S6 (phospho-Ser235/236, phospho-Ser240/244), which was accompanied by an upregulation of phospho-Ser473-Akt. However, TSC2-deficient tumor cells were less capable of regulating the phosphorylation level of these proteins upon T cell challenge (Fig. 3C). The same regulation was also observed in the A549 lung cancer cell line **(Fig. EV3B)**. This indicates that tumor cells respond to T cell challenge by redirecting their mTOR signaling toward a mTORC2-skewing phenotype, and that this regulation is impaired in *TSC2*-depleted tumor cells **(Fig. EV3C**). These results suggest an important role of TSC2 in regulating tumor tolerance to T cell attack via orchestrating the optimal ratio between mTORC1 and mTORC2 signaling.

Given these observations, we hypothesized that the shift towards mTORC1 signaling in TSC2-deficient tumor cells could account for the sensitization towards cytotoxic T cell attack, whereas inhibiting mTORC1 should reverse this phenotype. To test this, we treated T cell-challenged TSC2-deficient melanoma cells with LY2584702, an inhibitor of the S6 kinase, a crucial node for mTORC1 signal relay (Meyuhas, 2008; Magnuson *et al*, 2012). This caused TSC2-deficient cells to show a marked decrease of phosphorylated ribosomal protein S6 **(Fig. EV3D)**. In line with the reduction in mTORC1 signaling, when LY2584702 was added during T cell co-culture competition assay, the T cell sensitization induced by *TSC2* depletion was partially rescued (Fig. 3D). These results indicate that a TSC2-dependent orchestration of the mTORC1-mTORC2 signaling output contributes to modulating tumor cell sensitivity to T cells.

Whereas mTORC1 activation stimulates cell growth by upregulating multiple biosynthesis processes and inhibiting autophagy, its inhibition protects cells from inflammation-induced apoptosis and senescence (Kakiuchi *et al*, 2019). Furthermore, mTORC2 activation induces cell survival through Akt-mediated inhibition of apoptosis (Kennedy *et al*, 1997). In agreement with this, we found that *TSC2*-KO cells showed enhanced baseline apoptosis with a stronger expression of cleaved RIPK, caspase 3 and caspase 8, which was further enhanced upon T cell challenge (Fig. 3E). In addition to apoptosis, mTORC1 inhibition also protects cells from necroptosis (Abe *et al*, 2019), which serves as an alternative cell death signaling. By performing the competition assay in the presence of either a necroptosis specific inhibitor (Nec-1s), an apoptosis inhibitor (Q-VD-Oph) or both, we found that *TSC2*-depletion-induced-T cell sensitivity was abolished by blocking apoptosis signaling, which was partially rescued by necroptosis inhibition (Fig. 3F). This indicates that *TSC2*-KO cells are more vulnerable to both apoptosis-and necroptosis-induced cell death. Together, these results suggest that TSC2 protects tumor cells from T cell-induced cell cytotoxicity by orchestrating a pro-survival/anti-apoptotic mTOR response.

Because a TSC2 inhibitor is not available, we explored the potential translational value of modulating mTOR signaling for immunotherapy by other means. By treating tumor cells with a combination of a phosphatase inhibitor (tautomycin (TC)) and an Akt inhibitor (Triciribine (AktV)), we aimed to increase S6 phosphorylation (mTORC1 activation) and reduce Akt phosphorylation (mTORC2 inhibition). Indeed, the combination treatment led to simultaneous induction of S6 phosphorylation and potent suppression of Akt phosphorylation, producing a mTORC1-skewing phenotype similar to what was seen for *TSC2* depletion **(Fig. EV3E and EV3F).** Strikingly and in accordance with the *TSC2*-KO phenotype, pharmacological modulation of mTOR signaling was accompanied by an increased sensitivity to T cell killing; this was seen in several tumor cell lines (Fig. 3G). Taken together, these results indicate that tumor cells adapt to cytotoxic T cell attack by shifting the balance of mTOR signaling towards increased mTORC2 activity to circumvent apoptosis and necroptosis. By reversing this balance, either through genetic inactivation of *TSC2* or pharmacological modulation of mTOR signaling, tumor cell sensitivity to cytotoxic T cell attack can be augmented.

### *TSC2* depletion enhances TRAIL receptor expression and sensitizes melanoma cells to TRAIL-induced death

In response to antigenic stimulation, CD8^+^ T cells produce and secrete various effector cytokines that contribute to induction of cell death (Russell & Ley, 2002; Martínez-Lostao *et al*, 2015; Farhood *et al*, 2019). To dissect which of those are crucial in sensitizing *TSC2*-depleted tumor cells to T cells, we treated either Ctrl or *TSC2*-KO tumor cells with different cytokines secreted by T cells. We found that *TSC2*-KO cells were consistently more sensitive to TRAIL treatment (Fig. 4A **and Fig. EV4A and EV4B).** Of note, a sensitizing effect was also achieved by treatment with a human TRAIL monoclonal agonist antibody (Conatumumab) (Fig. 4B).

**Figure 4:**
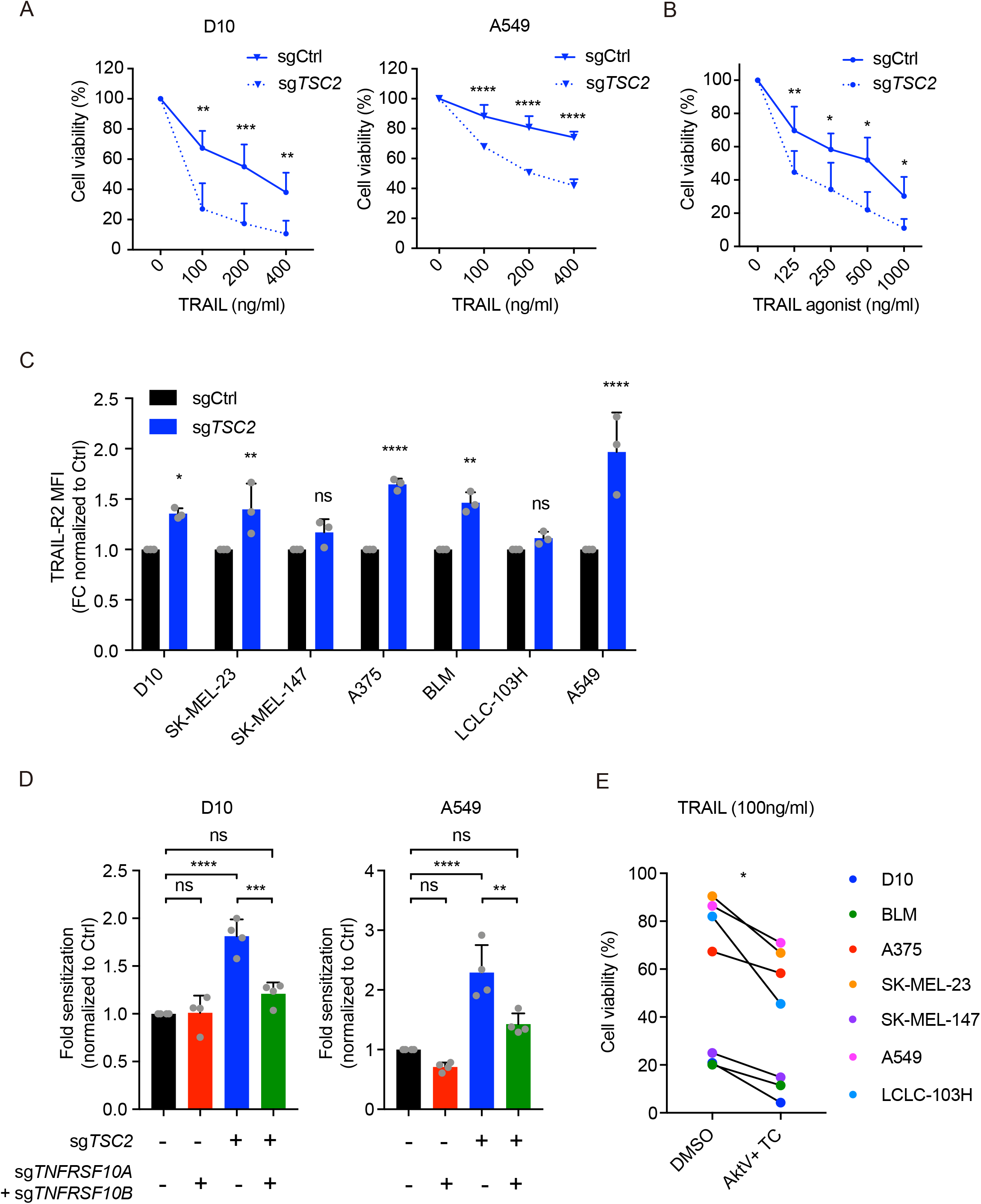
*TSC2* depletion enhances TRAIL receptor expression and sensitizes melanoma cells to TRAIL-induced death. A) Cell viability analysis by CellTiter-Blue assay of sgCtrl- or sgTSC2-expressing D10 melanoma and A549 lung cancer cells treated with TRAIL at indicated concentrations for three days. Statistical analysis was performed by two-tailed unpaired Student t test. For D10, error bars represent SD from three independent experiments (n = 3). For A549, error bars represent SD of pooled data from two independent experiments with three technical replicates each (n = 2). * P < 0.05; ** P < 0.01; *** P < 0.001; **** P < 0.0001. B) Cell viability analysis by CellTiter-Blue assay of sgCtrl- or sg*TSC2*-expressing D10 melanoma cells treated with TRAIL agonistic antibody Conatumumab, at indicated concentrations for two days. Statistical analysis was performed by two-tailed paired Student t test. Error bars indicate SD from three independent experiments (n = 3). * P < 0.05; ** P < 0.01; *** P < 0.001; **** P < 0.0001. C) Flowcytometry analysis of TRAIL-R2 surface expression level in multiple sg*TSC2-*expressing tumor cell lines compared to their sgCtrl-expressing controls. Statistical analysis was performed by two-way ANOVA with Sidak’s multiple comparisons test. Error bars represent SD of three independent experiments (n = 3). * P < 0.05; ** P < 0.01; *** P < 0.001; **** P < 0.0001. D) *In vitro* competition assay performed by mixing control and tumor cells with ablation for either *TSC2*, TRAIL receptors (*TNFRSF10A* and *TNFRSF10B*), or both in a co-culture with MART-1 T cells. Fold sensitization is normalized to Ctrl T cell-challenged groups. Statistical analysis was performed with a one-way ANOVA, followed by a Tukey post-hoc test. Error bars indicate SD of four independent experiments with different T cell donors. * P < 0.05; ** P < 0.01; *** P < 0.001; **** P < 0.0001. E) Cell viability analysis by CellTiter-Blue assay of TRAIL (100ng/ml) sensitivity in multiple parental melanoma and lung cancer cell lines in the presence of Tautomycin (TC, 150nM) and Akt Inhibitor V (AktV, 1.5uM) treatment, compared to DMSO-treated group. Statistical analysis was performed by Wilcoxon matched-pairs signed rank test of seven different cell lines. Representative graph of three independent experiments (n = 3). * P < 0.05; ** P < 0.01; *** P < 0.001; **** P < 0.0001.

TRAIL potently induces cell apoptosis by binding to the two death receptors, TRAIL-R1 (encoded by *TNFRSF10A*) and TRAIL-R2 (encoded by *TNFRSF10B*) (Wang & El-Deiry, 2003). To assess whether *TSC2* inactivation affects the expression of these receptors, we determined their expression on multiple Ctrl and *TSC2*-KO tumor cell lines by flow cytometry. This revealed markedly elevated levels of either TRAIL-R1, TRAIL-R2 or both (Fig. 4C **and Fig. EV4C)**. To evaluate the importance of TRAlL signaling in *TSC2*-depletion-induced-T cell sensitivity, competition assays were performed in the context of disrupted TRAIL signaling by knocking out both *TNFRSF10A* and *TNFRSF10B*. This perturbation diminished the sensitizing effect caused by *TSC2* depletion (Fig. 4D), establishing the requirement of TRAIL signaling in this setting. Moreover, pharmacologically manipulating mTOR signaling with TC/AktV combination treatment, which leads to a mTORC1-skewing phenotype, further sensitized parental tumor cells to TRAIL-induced cell death (Fig. 4E). These findings suggest a cross talk between mTOR regulation and TRAIL signaling, where *TSC2* may play a crucial role in the hub.

### Low TSC2 expression:TRAIL signaling ratio is associated with immune checkpoint blockade response in melanoma patients

Because *TSC2* depletion in tumor cells increases mTORC1 activity and elevates TRAIL receptor expression, which in turn sensitized tumor cells to T cell cytotoxicity, we next wished to explore any clinical implications. Examining the hallmark gene sets in The Cancer Genome Atlas (TCGA) analysis of melanoma, we found a negative correlation between *TSC2* expression and mTORC1 signaling, confirming the established function of TSC2 in mTOR regulation. Furthermore, *TSC2* expression level negatively correlated with that of TRAIL receptors (*TNFRSF10A* and *TNFRSF10B*), as well as of TRAIL signaling (Fig. 5A), in line with our findings above. Similar correlations were noticed among different cancer patient cohorts **(Fig. EV5A)**, indicating a more conserved role of TSC2 in TRAIL signaling regulation among different cancer types. To explore the influence of *TSC2* expression in cancer progression, we analyzed patient survival data from the TCGA melanoma cohort based on *TSC2* expression levels. Patients with melanomas expressing low levels of *TSC2* expression showed significantly prolonged survival **(Fig. EV5B)**. This association, while being subject to many factors including baseline immune pressure, is in agreement with our observation that TSC2 protects cells from apoptosis/necroptosis induced cell death.

**Figure 5:**
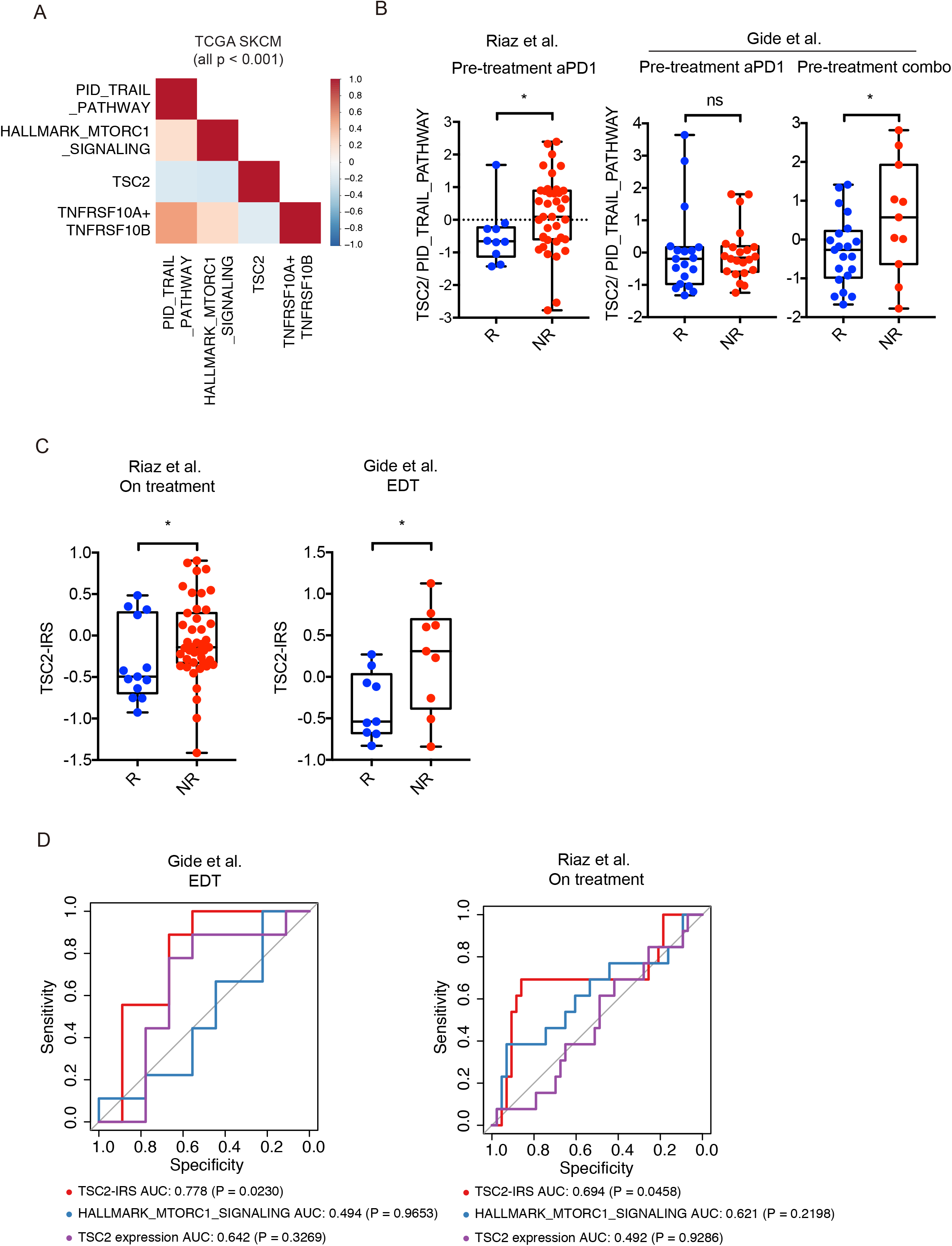
Low *TSC2* expression:TRAIL signaling ratio is associated with immune checkpoint blockade response in melanoma patients. A) Heat-map of Spearman correlation between *TSC2, TNFRSF10A/ TNFRSF10B* expression, mTORC1, and TRAIL signaling in TCGA SKCM (Skin Cutaneous Melanoma) baseline patient cohort. HALLMARK_MTORC1_SIGNALING (Liberzon *et al*, 2015), PID_TRAIL_PATHWAY (Schaefer *et al*, 2009) gene sets were taken from the Molecular Signatures Database (MSigDB). B) *TSC2* expression:TRAIL signaling ratio for predicting ICB treatment responses in pre-treatment patient samples (Riaz *et al*, 2017; Gide *et al*, 2019). Combo, anti-PD-1 plus anti-CTLA-4 combination treatment. Statistical significance was calculated with a Mann-Whitney test. * P < 0.05; ** P < 0.01; *** P < 0.001; **** P < 0.0001. C) TSC2 immune challenge signature expression in cohorts of patients after ICB treatment. Early during treatment (EDT), anti-PD-1 + anti-CTLA-4, and anti-PD-1 alone treatment cohorts. Statistical significance was calculated with a Mann-Whitney test. * P < 0.05; ** P < 0.01; *** P < 0.001; **** P < 0.0001. D) ROC curve analysis showing the probability of the indicated signatures as classifiers of ICB treatment response. Statistical significance between signatures and no predictive value (AUC=0.5) was calculated with bootstrapping. Partial response and complete response (PRCR) are defined as responder (R); stable disease (SD), progressive disease (PD) are defined as non-responder (NR).

Our findings indicate that upon TSC2 ablation, tumor cells induce their expression of TRAIL receptors, thereby increasing their susceptibility to T cell- or TRAIL-induced cell death. Therefore, we hypothesized that patients with lower *TSC2* expression (that is, with higher TRAIL receptor expression) together with a stronger baseline TRAIL signaling may be more sensitive to cancer immunotherapy and show better treatment outcome. To test this, we analyzed cohorts of melanoma patients before receiving ICB therapy. Indeed, we found that patients with a lower *TSC2* expression:TRAIL signaling ratio responded significantly better to ICB therapy (Fig. 5B) (Riaz *et al*, 2017; Gide *et al*, 2019). Of note, in the Gide data set, this ratio was significant only when patients were treated with both anti-PD-1 and anti-CTLA-4, likely triggering stronger immune pressure. These results indicate a correlation between the ratio of *TSC2* expression and TRAIL signaling and ICB treatment response of melanoma patients.

Lastly, in addition to *TSC2* single gene expression, we examined whether a TSC2 regulatory response upon immune challenge has any impact on ICB treatment outcome. We generated a TSC2 Immune Response Signature (TSC2-IRS) from differentially expressed genes between control and *TSC2*-depleted tumor cell lines under either T cell or TRAIL treatment. The TSC2-IRS was defined by the ratio of the differentially up-and down-regulated genes that overlapped between the two treatments **(Fig. EV5C)**. We analyzed the TSC2-IRS expression level in ICB-treated melanoma patient cohorts and found that responders showed a significantly lower TSC2-IRS expression (Fig. 5C). Moreover, TSC2-IRS expression significantly distinguished responding from non-responding patients, whereas *TSC2* single gene expression or mTORC1 signaling from the hallmark gene sets alone failed to do this (Fig. 5D). These clinical data point to a role for immune-induced TSC2 signaling in regulating tumor ICB response, supporting our functional data that *TSC2* inactivation augments tumor sensitivity toward immune pressure.

## Discussion

By re-interrogating the hits from a genome-wide CRISPR-Cas9 knockout screen we performed previously for critical determinants of tumor cell sensitivity to T cell killing (Vredevoogd *et al*, 2019), we identify here two negative mTOR regulators from the TSC complex, TSC1 and TSC2. Although these genes are established as tumor suppressors, their roles in determining tumor sensitivity to cytotoxic T cell challenge have not yet been described to our knowledge. We show here that TSC2 plays a dominant role over TSC1 in regulating tumor T cell sensitivity, and that this regulation is conducted, at least in part, through its control of mTORC1 and mTORC2 signaling balance; this regulation is impaired in *TSC2*-KO tumor cells upon T cell attack. We also demonstrate that by pharmacologically redirecting the mTOR downstream signal toward mTORC1 while inhibiting mTORC2 signaling (simultaneously sustaining phosphorylated ribosomal protein S6 and inhibiting Akt phosphorylation), tumor vulnerability to T cell killing can be induced (Fig. 6).

**Figure 6:**
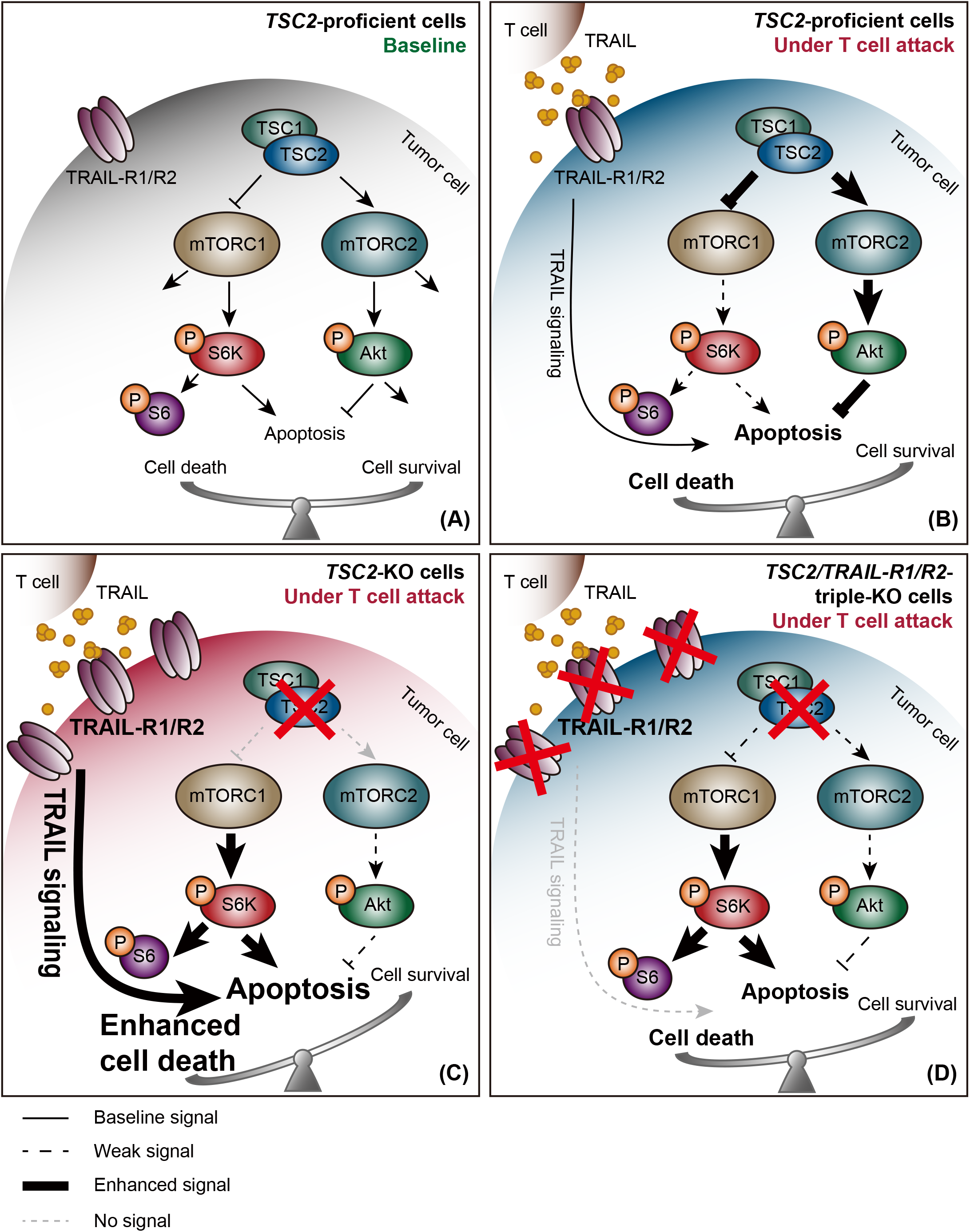
Modelling the role of TSC2 in governing tumor sensitivity to T cell killing. A) At baseline, Tuberous Sclerosis Complex (TSC1/2) suppresses mTORC1 signaling while inducing mTORC2 signaling. These factors, and their respective downstream targets, affect several pathways. In this study, we focused on the balance of tumor cell apoptosis and survival in the context of cytotoxic T cell attack. B) When challenged by cytotoxic T cells, tumor cells upregulate mTORC2 signaling and downregulate mTORC1 signaling, resulting in protection from cell death. C) Upon *TSC2* ablation, tumor cells (i) hyperactivate mTORC1 signaling, (ii) suppress mTORC2 signaling and (iii) elevate TRAIL receptor expression. Together, these events lead to increased susceptibility to T cell killing. D) When TRAIL-R1/R2 are ablated, *TSC2*-depleted tumor cells no longer receive extra TRAIL stimulation. As a consequence, only moderate cell killing is observed.

Cancer cells often show elevated mTORC1 signaling (Saxton & Sabatini, 2017; Grabiner *et al*, 2014; Sato *et al*, 2010; Gerlinger *et al*, 2012), resulting in enhanced cell growth associated with accelerated protein synthesis and metabolism (Düvel *et al*, 2010). However, several studies have shown that when encountering stress signals, cells downregulate mTORC1 signaling to lower their energy consumption rate and release the inhibition of autophagy, allowing for resource turnover (Ng *et al*, 2011; Aramburu *et al*, 2014). At the same time, they upregulate mTORC2 survival signal to inhibit cell apoptosis. Aligning with this, we found that tumor cells skew their mTOR signaling toward a mTORC2 phenotype when responding to stress induced by T cell challenge. This mTOR signaling regulation may allow tumor cells to balance their energy requirement and enhance apoptosis resistance to survive under unfavorable conditions.

Acting as the central regulator of both mTOR1 and mTORC2 signaling, the TSC1-TSC2 complex is considered to be a central integrator of external stress. It is essential for triggering proper stress responses through balancing the mTOR signaling level (Aramburu *et al*, 2014; Menon *et al*, 2014; Demetriades *et al*, 2014). TSC1-TSC2 complex inhibits mTORC1 signaling by regulating Rheb activity and activating mTORC2 signaling through direct interaction (Huang *et al*, 2008). It also plays a crucial role in the crosstalk between mTORC1 and mTORC2 signaling through PI3K/Akt feedback regulation. Thereby, Akt directly phosphorylates TSC2 to suppress the inhibitory function of TSC2 on Rheb and mTORC1, limiting TSC2’s inhibition of mTORC1 signaling (Manning et al, 2002; Huang & Manning, 2009; Cai et al, 2006; Potter et al, 2002). In agreement with our observations in multiple tumor cell lines, *TSC2* depletion interrupts the feedback regulation of mTORC2/Akt on mTORC1 signaling, leading to constitutively hyperactivated mTORC1 while suppressing mTORC2 signaling. This result confirms the status of TSC2 as a core regulator of the mTOR signaling balance.

TSC-deficient cells are more vulnerable to various cell death stimuli due to the impaired autophagy function caused by constitutive mTORC1 activation, while they are highly apoptotic due to diminished Akt signaling (Ng *et al*, 2011). In this study, we show that once *TSC2*-ablated tumor cells encounter cytotoxic T cell stress, they are less capable of downregulating mTORC1 signaling and upregulating mTORC2 signaling. As a result, *TSC2*-depleted cells continue to display a higher mTORC1/mTORC2 signaling ratio compared to TSC2-proficient tumor cells. Hyperactivated mTORC1 signaling is known to induce apoptosis owing to a constantly high metabolism rate and suppressed resource turnover from autophagy inhibition (Ng *et al*, 2011; Düvel *et al*, 2010). When treating *TSC2*-KO cells with LY2584702 (a S6 kinase inhibitor), we observed downregulation of mTORC1 signaling, which was associated with reduced T cell sensitivity. Together with the established inhibition by mTORC1 of autophagy, our data support the finding that autophagy inhibition sensitizes tumor cells to T cell killing (Lawson *et al*, 2020). On the other hand, *TSC2*-depletion directly inhibits mTORC2, thereby releasing the Akt-inhibited apoptosis (Kennedy *et al*, 1997), similar to what we observed in this study. Of note, we found Akt phosphorylation to be more heterogeneously regulated among different cancer cell lines. This may be caused by other regulators that phosphorylate Akt independently of TSC/mTORC2 signaling (Bozulic *et al*, 2008). Our study supports previous findings that TSC-null cells are extremely sensitive to multiple stress signals, such as DNA damage, ER stress, energy starvation and apoptosis (Kang *et al*, 2010; Wang *et al*, 2013). Our results are also in line with the finding in TSC and Lymphangioleiomyomatosis (LAM) animal models that treatment with anti-PD-1 antibody or the combination of anti-PD-1 and anti-CTLA4 antibodies leads to the suppression of *TSC2*-null tumor growth and induces tumor rejection (Liu *et al*, 2018).

Mechanistically, we demonstrate that *TSC2*-depleted tumor cells are highly susceptible to TRAIL-induced cell toxicity through its binding to death receptors TRAIL-R1 and TRAIL-R2. Specifically, we found that *TSC2*-depleted tumor cells elevate their expression of TRAIL receptors. Correspondingly, by blocking TRAIL signaling during T cell attack, increased T cell sensitivity of tumor cells upon *TSC2*-depletion could be completely rescued. Our finding indicates the existence of crosstalk between TSC2/mTOR signaling and TRAIL sensitivity, which supports previous studies that activation of the Akt survival pathway leads to TRAIL resistance in tumor cells (Chen *et al*, 2001; Nesterov *et al*, 2001).

Numerous mTOR inhibitors have been developed and are being tested for cancer treatment in the clinic (Chiarini *et al*, 2015; Hua *et al*, 2019). However, long-lasting anti-cancer effects are rare and patients often relapse due to various resistance mechanisms. Potential mechanisms have been suggested, including an activation of PI3K/Akt/mTORC2 survival signaling (Sun *et al*, 2005; Shi *et al*, 2005) and the upregulation of PD-L1 expression in cancer cells (Lastwika *et al*, 2016; Deng *et al*, 2019). On the other hand, mTOR inhibition is associated with immunosuppressive properties, since both mTORC1 and mTORC2 are required for proper T cell activation (Zheng *et al*, 2007; Colombetti *et al*, 2006) and trafficking (Sinclair *et al*, 2008). Moreover, mTORC1 and mTORC2 have distinct effects on fate decisions during immune cell differentiation (Chi, 2012; Delgoffe *et al*, 2011; Rao *et al*, 2010; Delgoffe *et al*, 2009). These findings emphasize the importance of uncoupling mTORC1 and mTORC2 signaling in any future cancer treatment.

TRAIL agonists have shown promising clinical benefit in cancer treatment (Snajdauf *et al*, 2021). Combination therapies are being explored, for example with mTOR inhibitors, aiming to boost the effect of TRAIL (Snajdauf *et al*, 2021). In this study, we show that induction of tumor cell death by *TSC2* depletion could be recapitulated by treatment with Conatumumab, a TRAIL agonist monoclonal antibody that is being been evaluated in several clinical trials. This opens up a potential translational value of targeting TSC2 in combination with TRAIL cancer therapy. Although the detailed mechanism of how TSC2/mTOR signaling regulates TRAIL receptor expression is to be explored, our finding provides in principle a rationale for selecting mTOR modulators as candidates for TRAIL combination therapy. Inhibiting mTOR signaling has a considerable impact on immune cell development and function (Dumont *et al*, 1990; Grolleau *et al*, 2002; Mills & Jameson, 2009). Therefore theoretically, targeting TSC2 in combination with TRAIL treatment, may bypass the undesired immunosuppressive effect of combining mTOR inhibitors with ICB therapy, which is highly dependent on immune cell functions.

Overall, our study uncovers crosstalk between TSC2 regulation and TRAIL signaling, and provides a novel concept for disrupting the mTORC1/mTORC2 balance to enhance tumor susceptibility to immune challenge. Further mechanistic study will be required to fully dissect the complexity of these signaling networks. This may allow us to identify specific targets for orchestrating an optimal mTORC1/2 signaling ratio in combination with TRAIL treatment in cancer therapy, aiming to avoid potential immunosuppressive effects.

## Materials and Methods

### Cell lines and primary cultures

D10, SK-MEL-23, SK-MEL-28, SK-MEL-147, A375, BLM, LCLC-103H and HEK293T cell lines were retrieved from the Peeper laboratory cell line stock. A549 cells were obtained from Wilbert Zwart. Human melanoma and lung cancer cell lines without endogenous HLA-A201 or MART-1 expression (SK-MEL-28, SK-MEL-147, A375, BLM, A549, LCLC-103H) were transduced with lentiviral constructs encoding both components. The B16-F10 cell line was obtained from ATCC and was lentivirus-transduced to express the full-length ovalbumin (OVA) protein. OVA-expressing cells were selected with hygromycin (250 µg/ml, 10687010, Life Technologies). All cell lines were cultured in DMEM (GIBCO), supplied with 10% fetal bovine serum (Sigma) and 100U/ml of Penicillin-Streptomycin (GIBCO). All cell lines were regularly tested for mycoplasma by PCR (Young *et al*, 2010) and were authenticated using the STR profiling kit from Promega (B9510).

### Isolation, generation and maintenance of MART-1 TCR CD8 T cells

MART-1 TCR CD8 T cells were generated as previously described (Vredevoogd *et al*, 2019). Briefly, primary CD8 T cells were isolated from fresh, healthy male or female donor buffycoats (Sanquin, Amsterdam, the Netherlands), activated for 48 hours in human CD8 T cell media (RPMI Medium (GIBCO) containing 10% human serum (H3667, Sigma-Aldrich), 100U/ml of Penicillin-Streptomycin, 100U/ml IL-2 (Proleukin, Novartis), 10ng/ml IL-7 (11340077, ImmunoTools) and 10ng/ml IL-15 (11340157, ImmunoTools)) with plate-coated αCD3 and αCD28 antibodies (16-0037-85 and 16-0289-85, eBioscience) and spinfected with MART-1 TCR retrovirus on Retronectin coated (TB T100B, Takara) non-tissue culture treated plates. Cells were harvested and maintained in human CD8 T cell media 24h after transduction. One week after retroviral transduction, MART-1 TCR expression was confirmed by flow cytometry (α-mouse TCR β chain, 553172, BD Pharmingen) and cells were cultured in RPMI containing 10% fetal bovine serum (Sigma), 100U/ml of Penicillin-Streptomycin (GIBCO) and 100U/ ml IL-2 (Proleukin, Novartis).

### Knockout and overexpression cell line generation

Knocking out and overexpressing genes of interest in different cell lines were done by lentiviral transduction. For gene knockouts, sgRNAs were cloned into lentiCRISPR-v2 (#52961, Addgene) plasmid using a SAM target sgRNA cloning protocol (S. Konermann, Zhang lab, 2014). For *TSC2* reconstitution, full length *TSC2* cDNA containing a CRISPR-Cas9-resistant silent mutation was cloned into pCDH-blast plasmid. Lentivirus was produced by transfecting HEK293T cells with psPAX2 (#12260, Addgene) and pMD2.G (#12259, Addgene) using polyethylenimine. The media was refreshed with OptiMEM (31985062, GIBCO) containing 2% fetal bovine serum 24 h after transfection. Supernatant was harvested 72 h post-transfection, filtered and stored at −80°C. Tumor cells were transduced with lentivirus, together with polybrene (8 μg/ml), and cell media was refreshed 24h later. Cells were selected with antibiotics for at least one week. *TSC1*/*TSC2* double knockout cells or *TSC2* reconstitution in knockout cells were done sequentially by using both puromycin-selectable and blasticidin-selectable plasmids. *TNFRSF10A /TNFRSF10B* double knockout cells were purified by cell sorting after antibiotic selection.

sgRNA targeting sequences:

sg_hCtrl: 5’-GGTTGCTGTGACGAACGGGG -3’

sg_h*TSC1*: 5’- CGAGATAGACTTCCGCCACG -3’

sg_h*TSC2*-1: 5’- CAGAGGGTAACGATGAACAG -3’

sg_h*TSC2*-2: 5’- TCCTTGCGATGTACTCGTCG -3’

sg_h*TSC2*-3: 5’- ATTGTGTCTCGCAGCTGATG -3’

sg_h*TNFRSF10A*: 5’- AGCCTGTAACCGGTGCACAG -3’

sg_h*TNFRSF10B*: 5’- AGGTGGACACAATCCCTCTG -3’

sg_mCtrl: 5’- AAAAAGTCCGCGATTACGTC -3’

sg_m*Tsc2*-1: 5’- TCATTCGGATGCGATTGTTG -3’

sg_m*Tsc2*-2: 5’- AGTTCTTGAGAGAGTAGAGC -3’

sg_m*Tsc2*-3: 5’- GGTCAGCAGGTCATGGACGA -3’

### *In vitro* competition assay

Parental or gene-modified cells were stained with either the CellTrace CFSE Cell Proliferation Kit (C34554, CFSE; Thermo Scientific) or the CellTrace Violet Cell Proliferation Kit (C34557, CTV; Thermo Scientific) following manufacturer’s instructions. Stained cells were mixed at a 1:1 ratio and challenged with either MART-1 T cells or Ctrl T cells for 3 days. For drug treatment, indicated compounds were added in the media together with T cell coculture, LY2584702 (S7704, SelleckChem), Nec-1s (50-429-70001, Sigma-Aldrich), Q-VD-Oph (S7311, SelleckChem). For TRAIL treatment competition assay, 100ng/ml sTRAIL/Apo2L (310-04, Peprotech) was added to the culture media. The percentage CFSE-and CTV-positive cells was analyzed by flow cytometry. Sensitivity was calculated by the ratio of control cells to gene-modified cells under MART-1 T cells challenge normalized to the Ctrl T cell condition.

### *In vitro* cytotoxicity assay

1×10^4^ tumor cells were seeded per well into 96-well culture plates (Greiner). Recombinant Human IFNα-1b (11343594, ImmunoTools), IFNβ-1b (11343543, ImmunoTools), IFNγ (Peprotech), TNFα (300-01A, Peprotech), TNFβ (300-01B, Peprotech), sFas Ligand (310-03H, Peprotech), sTRAIL/Apo2L (310-04, Peprotech), Tautomycin (580551, Sigma-Aldrich), Triciribine (Akt Inhibitor V, 124012, Merckmillipore), Conatumumab (TAB-203, Creative Biolabs Inc.) were added at indicated concentrations. Cells were incubated for 3 days before viability analysis. Cell viability was read using Cell Titer Blue Viability Assay (G8081, Promega) according to manufacturer’s instruction.

### *In vivo* competition assay and mouse model

D10 cells were first lentivirally transduced with sgCtrl or sg*TSC2* and selected with puromycin for one week as described above. Then, cells were lentivirally transduced with eGFP (pLX304-EGFP-Blast) or mCherry (pLX304-mCherry-Blast) expression plasmids and sorted. Cells were mixed at 1:1 ratio prior to injection. 1×10^6^ mixed cells per mouse were subcutaneously injected into immune-deficient NSG-B2m mice (n=10, The Jackson Laboratory, Strain #:010636) with Matrigel (354230, Corning). Tumor growth was monitored three times per week. Mice were randomized 12 days after tumor injection, and either 5 x 10^6^ MART-1 or Ctrl human CD8 T cells were intravenously injected into the tail vein, followed by daily 100.000 U IL-2 (Proleukin, Novartis) intraperitoneal injection for 3 consecutive days. Researchers were blinded for treatment given. Tumors were harvested 8 days post ACT and digested into single cell suspensions. EGFP- and mCherry-positive cells were analyzed by flow cytometry. Mice without tumor outgrowth or failed to receive proper ACT were excluded.

### Flow cytometry

Cells were harvested and stained with fluorescent-conjugated antibodies targeting surface molecules of interest according to manufacturer’s protocol. Samples were analyzed with Fortessa flow cytometer (BD Bioscience). Antibodies against human CD261 (307207, Biolegend), CD262 (307405, Biolegend), and Live/Dead Fixable Near-IR Dead Cell Stain Kit (L34976, Thermo) were used.

### Immunoblotting

Cells were washed with PBS, scrape-harvested, and lysed for 30min on ice with RIPA buffer (50mM TRIS pH 8.0, 150mM NaCl, 1% Nonidet P40, 0.5% sodium deoxycholate, 0.1% SDS) supplemented with Halt Protease and Phosphatase inhibitor cocktail (78444, Fisher Scientific) for phosphoprotein blotting. Samples were centrifugated at 17000g, supernatant was collected and protein concentration was measured by Bradford Protein Assay (500-0006, Bio-Rad). To prepare immunoblot samples, protein concentration was normalized and 4xLDS sample buffer (15484379, Fisher Scientific) containing 10% β-Mercaptoethanol (final concentration 2.5%) was added, following by 5 min incubation at 95°C. Samples were size-separated on 4–12% NuPAGE Bis-Tris polyacrylamide-SDS gels (Invitrogen) and transferred on nitrocellulose membranes (IB301031, Invitrogen, iBlot™ Transfer Stack). Blots were blocked in 4% milk powder in 0.2% Tween/PBS (PBST) and incubated at 4°C overnight with primary antibodies. After washing by PBST, secondary antibodies were applied for 1h at room temperature. Blots were then washed by PBST and developed with SuperSignal West Dura Extended Duration Substrate (34075, Thermo Scientific) and luminescence was captured Luminescence signal was detected by either Amersham Hyperfilm high performance autoradiography film or Bio-Rad ChemiDoc imaging system with default settings. Primary antibodies against TSC1 (6935, Cell Signaling Technology), TSC2 (4308, Cell Signaling Technology), Akt (sc-8312, Santa Cruz Biotechnology), pAktSer473 (4060, Cell Signaling Technology), S6R (2217, Cell Signaling Technology), pS6Ser240/244 (2215, Cell Signaling Technology), pS6Ser235/236 (2211, Cell Signaling Technology), Caspase 3 (9665, Cell Signaling Technology), cleaved Caspase 3 (9664, Cell Signaling Technology), Caspase 8 (4790, Cell Signaling Technology), cleaved Caspase 8 (9748, Cell Signaling Technology), RIPK1 (3493, Cell Signaling Technology), Vinculin (4650, Cell Signaling Technology), α-Tubulin (T9026, Sigma), HSP90 (sc-7947, Santa Cruz Biotechnology), cyclophilin B (43603, Cell Signaling Technology) were used. Horseradish peroxidase-conjugated secondary antibodies against mouse IgG (G21040, Thermo Scientific), rabbit IgG (G21234, Invitrogen) were used.

### Sample preparation for generating TSC2 immune challenge signature

D10 melanoma and A549 lung cancer cell lines expressing sgCtrl or sg*TSC2* were seeded into 10cm tissue culture dishes at 70% for 48h. Tumor cells were challenged with CFSE (423801, Biolegend) pre-labelled MART-1 T cells, with T cell to tumor ratio causing 50% tumor cell killing, or sTRAIL/Apo2L (310-04, Peprotech) (D10: 10ng/ml; A549: 100ng/ml) overnight. Supernatant containing cell debris and T cells was discarded and attached cells were harvested by trypsinization. Samples were washed with PBS and stained with DAPI. Pure viable tumor cells were sorted by FACSAria Fusion Cell Sorters gating on DAPI-, CFSE-populations and sent for RNA sequencing.

### Whole-genome CRISPR-KO screen data analysis

Count data from the whole genome screen (Vredevoogd *et al*, 2019) was re-analyzed using MAGeCK (v0.5.7) using the second best sgRNA method (Li *et al*, 2014). To make this analysis more robust, sgRNAs with low read counts (<50) were filtered from this analysis.

### Data Resources and bioinformatic analysis

Raw gene counts of the TCGA SKCM data set were downloaded with TCGAbiolinks in R (4.0.2) from the GDC Data Portal together with metadata. Read count data was pre-processed and normalized using DESeq2 (1.30.0). Expression data of the pan-cancer TCGA cohorts was obtained using query option on the cBioPortal website. Survival analysis was performed on the overall survival data in months and the observed overall survival in months (Xiong *et al*, 2019), using the top and bottom quantiles of the *TSC2* expression for grouping the samples.

For the anti-PD-1-treated patient cohort (Riaz *et al*, 2017), the raw counts were downloaded from NCBI’s GEO (GSE91061). For a second patient cohort, containing patients treated with either anti-PD-1 monotherapy or combined anti-PD-1 and anti-CTLA-4 (Gide *et al*, 2019), the RNA sequencing data was downloaded from the European Nucleotide Archive (ENA) under project PRJEB23709. The fastq files were mapped using STAR (2.6.0c) with default settings on two-pass mode. The raw counts were generated using HTSeq (0.10.0). For both cohorts, the raw read count data were pre-processed and normalized using DESeq2 (1.30.0). Z-scores were obtained from the normalized read counts by subtracting the row means and scaling by dividing the columns by the SD.

For the data on the control and *TSC2*-depleted cell lines after cytotoxic T cell or TRAIL challenge used for generating the TSC2 immune response signature (TSC2-IRS), the fastq files were mapped to the GRCh38 human reference genome (Homo.sapiens.GRCh38.v82) using STAR (2.7.3a) with default settings on two-pass mode. Count data was generated with HTSeq (0.12.4) and pre-processed and normalized using DESeq2 (1.30.0). The TSC2-IRS signature was generated by overlapping the up and down differentially expressed genes (DEGs) between the *TSC2*-depleted cell lines versus the control of both T cells and TRAIL treatments. Significant DEGs were defined by an adjusted p-value of less than 0.01 and a minimum fold change (fc) of 0.15 or maximum fc of −0.15, for genes that were either up or down in *TSC2*-depleted cell lines respectively. For applying the TSC2-IRS signature on the clinical data, we took the ratio between the down and up signatures.

### Statistics

When comparing two groups, a Two-tailed Student t test was performed for normally distributed data, or by two-tailed Mann-Whitney test with Bonferroni correction for data that was not normally distributed. When comparing more than one group of data to one control group, one-way ANOVA with Holm-Sidak’s multiple comparisons test was performed when data is normally distributed, or Kruskal-Wallis test with Dunn’s post hoc test was used when data was not normally distributed. Tukey’s post-hoc analysis was used for multiple comparisons between all groups. Data distribution normality was analyzed by Shapiro-Wilk test. All analyses were performed by Prism (Graphpad Software Inc.). P value lower than 0.05 was defined as statistically significant. For *in vivo* experiments, sample size estimation for experimental study design was calculated by G*Power (Faul *et al*, 2007).

### Data availability

The RNA-Seq dataset used in this study to generate TSC2-IRS is available in the Gene Expression Omnibus (GEO) database (GSE201514), and assigned the links as below: https://www.ncbi.nlm.nih.gov/geo/query/acc.cgi?acc=GSE201514 (access upon request).

### Animal welfare

All animal studies were approved by the animal ethics committee of the Netherlands Cancer Institute (NKI) and performed under approved NKI CCD (Centrale Commissie Dierproeven) projects according to the ethical and procedural guidelines established by the NKI and Dutch legislation. Male or female NSG-B2m (The Jackson Laboratory) mice were used after at 8 weeks or older. Mice were housed in one-time use standard cages at controlled air humidity (55%), temperature (21 °C) and light cycle. All housing material, food and water were autoclaved or irradiated before use.

## Acknowledgments

We thank all members in the Peeper laboratory, especially G. Apriamashvili, B. de Bruijn, O. Krijgsman, and T. Kuilman, and our colleagues in the Division of Molecular Oncology and Immunology for their valuable input. We thank T. Schumacher and W. Zwart for sharing cell lines and input R. Bernards for sharing Conatumumab. Moreover, we would like to thank The Netherlands Cancer Institute Flow Cytometry facility, Genomic Core facility, Animal Laboratory for their contributions. This work was supported by Oncode Institute.

## Author Contributions

Chun-Pu Lin: Conceptualization, Data curation, Investigation, Methodology, Project administration, Validation, Visualization, Writing – original draft. Joleen Traets: Formal analysis, Software. David W. Vredevoogd: Conceptualization, Formal analysis, Methodology. Nils L. Visser: Resources. Daniel S. Peeper: Conceptualization, Funding acquisition, Supervision, Writing – review & editing.

In addition to the above CRediT contributor roles taxonomy, the detail contribution descriptions are:

C.L. and D.S.P. designed, administrated and performed all experiments. J.T. performed bioinformatics analyses. D.W.V. re-analyzed the screen and identified TSC2. C.L., D.W.V. and D.S.P. discussed the data. D.W.V. and N.L.V. provided technical support. C.L. and D.S.P. wrote the manuscript. All authors revised and approved the manuscript. The project was supervised by D.S.P.

## Conflict of interest

D.S.P. is a co-founder, shareholder and advisor of Immagene, which is unrelated to this study. The other authors declare that they have no conflict of interest.

**Figure EV1: Whole-genome CRISPR-Cas9 knockout screen identifies TSC2 as a negative regulator of tumor sensitivity to CTL killing.**

A) *In vitro* tumor T cell co-culture competition assay in D10 human melanoma cells expressing different sgRNAs targeting human *TSC2*. Error bars represent SD of five independent experiments with different T cell donors (n = 5). Statistical analysis was performed by one-way ANOVA with Holm-Sidak’s multiple comparisons test. * P < 0.05; ** P < 0.01; *** P < 0.001; **** P < 0.0001.

B) Western blot analysis of *TSC2* expression in cells used in (A), (n = 2).

C) *In vitro* tumor T cell co-culture competition assay in B16-F10-OVA murine melanoma cells expressing different sgRNAs targeting murine *Tsc2*. Error bars represent SD of pooled data from two independent experiments with three technical replicates each (n = 2). Statistical analysis was performed by Kruskal-Wallis test with Dunn’s post hoc test. * P < 0.05; ** P < 0.01; *** P < 0.001; **** P < 0.0001.

D) Western blot analysis of *TSC2* expression in cells used in (C), (n = 2).

**Figure EV2: TSC2 ablation sensitizes melanoma cells to CTL killing in an *in vivo* tumor ACT model.**

Repeat of *in vivo* competition assay from Figure 2, with reversed fluorescent color labeling.

A) Tumor growth upon adoptive cell transfer with Ctrl (n = 10) or MART-1 (n = 10) T cells. Data points represent average tumor volume, and error bars represent SEM.

B) Flow cytometry plot of both tumor mix input and representative output of the *in vivo* competition assay.

C) Quantification of the *in vivo* competition assay (output) at end point by flow cytometry analysis. Error bars indicate SD. Statistical analysis was performed by Mann–Whitney test. * P < 0.05; ** P < 0.01; *** P < 0.001; **** P < 0.0001.

**Figure EV3:**

***TSC2* depletion-induced deregulation of mTOR signaling increases tumor cell death by CTL attack.**

A) Western blot analysis of baseline mTOR signaling in melanoma cells expressing sgCtrl, sg*TSC1*, sg*TSC2* or both. All blots, except D10, were run in parallel and serve to make comparisons within individual cell lines. SK-MEL-23 lysate was run in parallel with other cell lines but on a separate gel.

B) Western blot analysis of mTOR signaling in sgCtrl- or sg*TSC2*-expressing A549 lung cancer cells upon MART-1 T cell challenge for indicated time. Representative of 2 experiments with different T cell donors (n = 2).

C) Western blot quantification for mTOR1/mTORC2 signaling ratio changes of sgCtrl- or sg*TSC2*-expressing D10 melanoma cells upon T cell challenge, as shown in Figure 3C. Signaling ratio was calculated as in Figure 3B. Error bars represent SEM of the quantification from three independent experiments with different T cell donors (n = 3).

D) Western blot analysis of mTOR signaling in sgCtrl- or sg*TSC2*-expressing D10 cells upon DMSO or LY2584702 (5uM) treatment for indicated time. Representative of 3 experiments (n = 3).

E) Western blot analysis of mTOR signaling in parental D10 melanoma cells upon DMSO or Tautomycin (150nM) and Akt Inhibitor V (1.5uM) combination treatment for indicated time. Representative of 3 experiments (n = 3).

F) Western blot quantification for mTOR1/mTORC2 signaling ratio changes in E). Error bars represent SEM of the quantification from three independent experiments (n = 3).

**Figure EV4: *TSC2* depletion enhances TRAIL receptor expression and sensitizes melanoma cells to TRAIL-induced death.**

A) Cell viability analysis by CellTiter-Blue assay of sgCtrl- or sg*Tsc2*-expressing D10 melanoma (n = 3) and A549 lung cancer (n = 2) cells treated with different cytokines at indicated concentrations for three days. Representative graphs of three and two independent experiments, respectively. Error bars represent SD of three technical replicates.

B) *In vitro* competition assay of sgCtrl- and sg*TSC2*-expressing D10 melanoma or A549 lung cancer cell lines exposed to TRAIL (50ng/ml) treatment, normalized to untreated control groups. Statistical analysis was performed by two-tailed unpaired Student t test. Error bars represent SD of four independent experiments (n = 4). * P < 0.05; ** P < 0.01; *** P < 0.001; **** P < 0.0001.

C) Flowcytometry analysis of baseline TRAIL-R1 surface expression level in multiple sg*TSC2*-expressing tumor cell lines comparing to their sgCtrl-expressing controls. Statistical analysis was performed by two-way ANOVA with Sidak’s multiple comparisons test. Error bars represent SD of three independent experiments (n = 3). * P < 0.05; ** P < 0.01; *** P < 0.001; **** P < 0.0001. *), non-detectable as MFI close to background staining.

**Figure EV5: Low *TSC2* expression:TRAIL signaling ratio is associated with immune checkpoint blockade response in melanoma patients.**

A) Heat-map of Spearman correlation between expression of *TSC2* and *TNFRSF10A* and *TNFRSF10B*, and mTORC1 or TRAIL signaling in multiple baseline TCGA cancer patient cohorts.

B) Kaplan-Meier survival curves of TCGA SKCM patient cohort with high (top 25%) or low (bottom 25%) *TSC2* expression. Significance between curves was calculated with a regular log-rank test.

C) Heatmap of overlapping differentially expressed genes between *TSC2*-depleted and control cells in D10 and A549 cell lines upon either MART-1 T cell or TRAIL treatment.

